# The intracellular Ca^2+^ sensitivity of transmitter release from neocortical boutons

**DOI:** 10.1101/2024.02.26.582057

**Authors:** Grit Bornschein, Simone Brachtendorf, Abdelmoneim Eshra, Robert Kraft, Jens Eilers, Stefan Hallermann, Hartmut Schmidt

**Affiliations:** Carl-Ludwig-Institute for Physiology, Medical Faculty, Leipzig University, Germany

## Abstract

Synaptotagmin 1 (Syt1) and Syt2 are the main Ca^2+^ sensors triggering synchronous release in the brain. The Ca^2+^-sensitivity of Syt2-triggered release has been studied in detail. However, for Syt1, the dominating isoform in the neocortex, quantitative detail is lacking. We measured the Ca^2+^-dependency of Syt1-triggered release at layer 5 pyramidal neuron synapses by laser photolysis of caged Ca^2+^. Syt1-triggered release had high Ca^2+^ affinity and positive cooperativity (EC_50_, 20 μM; Hill coefficient, 3.57). It was steep in a dynamic range between ∼10 and ∼30 μM that was covered by action potential-evoked release. A kinetic model reveals significant differences to models of Syt2-triggered release. Our results suggest that Syt1 optimizes neocortical synapses for high reliability at moderate local Ca^2+^ elevations and for high plastic controllability.

---

Fast synchronous release of neurotransmitters from presynaptic boutons is initiated by action potentials (APs) opening voltage-gated Ca^2+^ channels. The inflowing Ca^2+^ binds to a sensor protein, which then triggers the release process (*1*). Due to the steepness and short duration of the Ca^2+^ gradient, chemical equilibrium is not established in this process. This makes the physical coupling distance between the Ca^2+^ channels and the sensor protein as well as the intracellular Ca^2+^-binding kinetics of the sensor central in the control of speed, reliability and plasticity of release (*2-7*).

While coupling distances have been studied in detail at different synapses in recent years (*8-14*), the intracellular Ca^2+^-binding kinetics have been less intensively investigated. Syt1 and Syt2 are the main Ca^2+^ sensors for synchronous release in the brain (*1, 15*). Syt2 is the dominating sensor in hindbrain synapses, while in forebrain synapses, particularly in the neocortex, the sensor is Syt1 (*15-18*). At the giant calyx of Held, the synaptic Ca^2+^-binding kinetics for Syt2-triggered glutamate release were quantified from presynaptic Ca^2+^ elevations and transmitter release rates in detail, resulting in a kinetic model of Ca^2+^-binding and release with five cooperative Ca^2+^-binding sites (*2, 3, 19*). In subsequent work, this ‘Syt2-model’ has been extended to account for release at low elevations of the presynaptic Ca^2+^ concentration (Δ[Ca^2+^]_i_) (*20*) relevant for asynchronous release (*18*), for developmental alterations (*21, 22*), for Syt2-triggered GABA release (*6*), and for priming and synaptic plasticity (*6, 23-25*).

At present, corresponding quantitative detail is lacking for the Ca^2+^-binding kinetics of Syt1-triggered release. Due to their small size and axo-dendritic location neocortical synapses provide less favorable experimental conditions than giant (*2, 3, 25*) or axo-somatic (*6*) Syt2-expressing synapses. In particular, the axo-dendritic locations of the small boutons pose a substantial hurdle for the identification of the connected boutons in intact brain tissue. Here, we have succeeded in solving these experimental difficulties and quantified the Ca^2+^-dependency of Syt1-triggered release at synapses connecting layer 5 pyramidal neurons (L5PNs).

## Identifying boutons connecting L5PNs with UV-laser photolysis

We combined UV-laser-based Ca^2+^ uncaging with quantitative green over red (G/R) two-photon Ca^2+^ imaging at synapses between L5PNs in the primary somatosensory cortex (S1) (**Fig. 1, Supplementary Fig. 1**; see **Methods**). After a paired recording had been established (*11*), presynaptic neurons were loaded via the somatic whole-cell patch pipettes with the caged Ca^2+^-compound DM-nitrophen (DMn), the green-fluorescent, low-affinity Ca^2+^ indicator dye Oregon Green 488 BAPTA-5N (OGB5N; *K*_D_, 22 μM) and the Ca^2+^-insensitive, red-fluorescent dye Atto-594. The typical synaptic connection between L5PNs is formed by 4-8 boutons distributed essentially over the basal dendrites. The boutons are situated close to the soma of the postsynaptic cell (<100 μm) and well separated from each other (≥100 μm) (*26*). To identify the connecting boutons among the multitude of boutons on an axon collateral, we performed ‘axonal walking’: Using the bright red fluorescence, we followed axon collaterals in the direction of the soma of the postsynaptic neuron and applied UV-flashes to boutons at distances ≤100 μm from the soma until photolysis induced both, a presynaptic Ca^2+^ signal and a flash-induced excitatory postsynaptic current (fEPSC; **Fig. 1**). Subsequently, brief UV-laser flashes of different intensity and duration were applied to yield different Δ[Ca^2+^]_i_. The Δ[Ca^2+^]_i_ were monitored at high temporal resolution by point-mode recordings from individual boutons. The diameter of our uncaging spot was ∼10 μm, i.e. 10 times smaller than the typical 100 μm or more spacing between boutons forming the synaptic connection (*26*). Hence, it is most likely that we measured mainly fEPSCs resulting from release from single boutons.

**Fig. 1.**
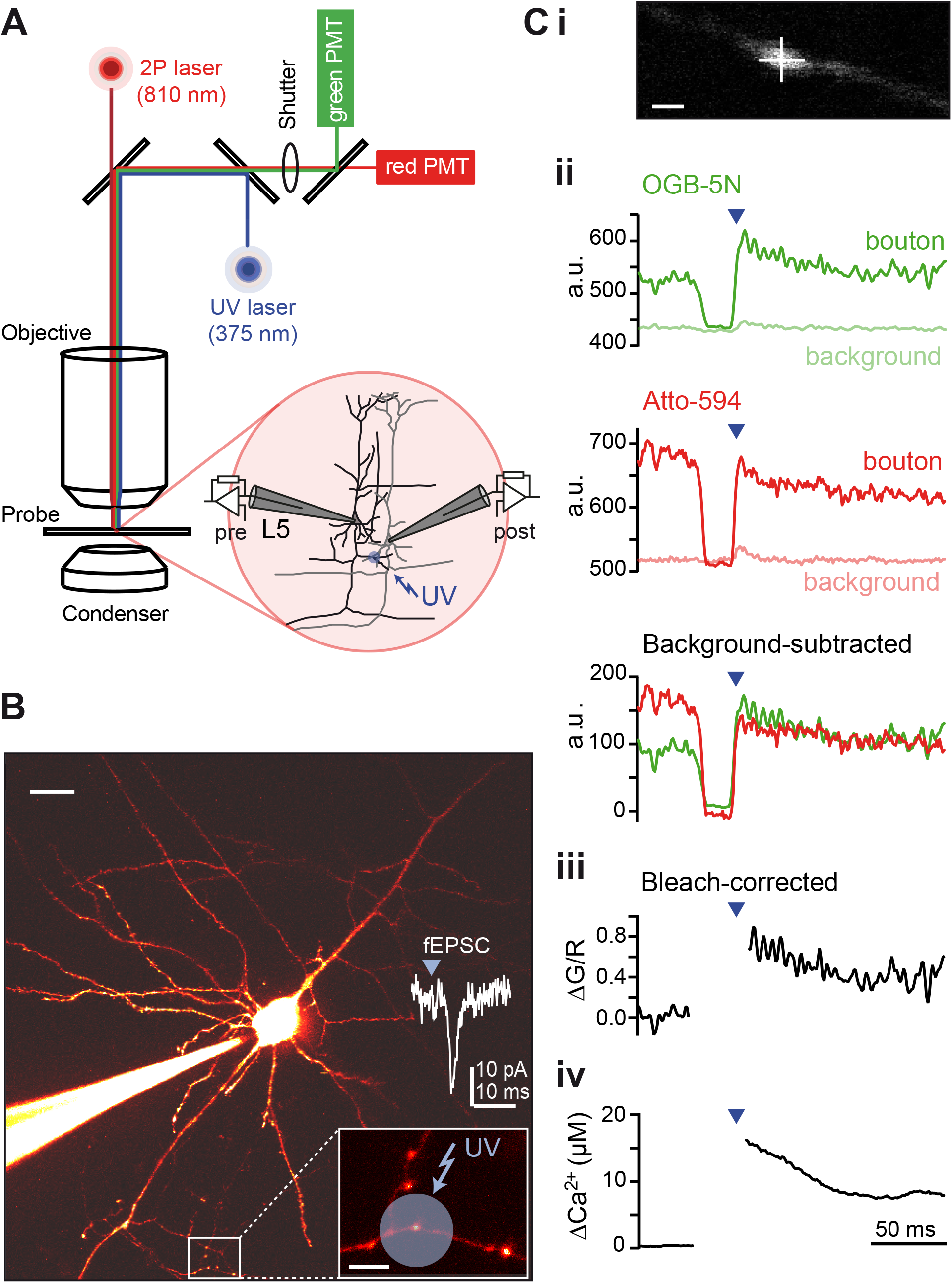
UV-photolysis induced Δ[Ca^2+^]_i_ and fEPSCs at pairs of connected L5PNs. **(A)** Scheme of the experimental setup showing the light path of the two-photon laser excitation (dark red), the UV laser illumination (blue), the emitted red and green fluorescence detected by gate-able photomultiplier tubes (PMTs, red and green respectively). Inset: Illustration of connected L5PNs that were identified by paired recordings. **(B)** Two-photon image of a presynaptic L5PN (scale bar 20 μm) and its axonal boutons (inset, scale bar 5 μm). The presynaptic neuron was loaded with OGB5N, Atto594, and DMn/Ca^2+^ via the patch pipette; Ca^2+^ elevations were induced by application of brief UV flashes to presumed presynaptic boutons until corresponding fEPSCs (inset) were recorded from the postsynaptic neurons (not labeled). (**Ci**) Two-photon image of a bouton located on an axon collateral from which flash-induced green (G) and red (R) fluorescence were recorded in the point-mode (cross, scale bar 1 μm). (**Cii**) Two-photon point mode recordings showing the green (G, *top*) and red (R, *middle*) fluorescence as detected by the two PMTs and following background subtraction (*bottom*; time of the UV flash indicated by blue arrow). The background subtraction eliminated a residual flash-induced artifact. (**Ciii**) Calculated ΔG/R signals with correction for bleaching in the red fluorescence (see **Methods**). (**Civ**) Δ[Ca^2+^]_i_ calculated from the ΔG/R signal based on a cuvette calibration.

Occasionally the UV spot covered more than one bouton in the x-y plane. Hence, we probed for homogeneity in UV-induced Δ[Ca^2+^]_i_ between boutons within the photolysis-area, using line-scan recordings that simultaneously assessed the fluorescence changes in these boutons. We found that ΔG/R were rather homogeneous with an average deviation of 11% between boutons (**Supplementary Fig. 2**; mean SD=11%, n=9 ROIs from 7 cells). Thus, although we may not always have imaged the connected bouton that evoked the fEPSCs through its release in the point mode, these data suggest that the corresponding error in the Δ[Ca^2+^]_i_ estimate will be moderate. These results indicate a reliable correlation between the Δ[Ca^2+^]_i_ measured at presynaptic boutons and the fEPSCs.

### Dose-response curve for the dependency of release on presynaptic Ca^2+^

The transmitter release activity was monitored by recording fEPSCs (**Fig. 1, 2A**). Since EPSC amplitudes are not a valid measure of presynaptic release rates (*2*), we obtained presynaptic release rates from multiquantal fEPSCs by the deconvolution method (*2, 27*), using an average quantal EPSC (qEPSC) evoked by APs in low [Ca^2+^]_e_ (*11*). The qEPSCs had an average amplitude of 6 pA (5-7 pA, n=7) and decayed monoexponentially with τ of 2.1 ms (1.5-4.0 ms; **Supplementary Fig. 3A**). Due to the small size of the fEPSCs, the resulting release rates were rather noisy. To reduce the noise, we integrated the release rates over time, which resulted in smooth cumulative release curves. The cumulative curves were typically monoexponential for fEPSCs evoked by small to intermediate Δ[Ca^2+^]_i_. They occasionally became biexponential for large presynaptic Δ[Ca^2+^]_i_ (> ∼18-20 μM; **Fig. 2B**). In the slow phase of the biexponential curves individual steps were frequently visible that may indicate fusion of few or individual vesicles. The cumulative curves were fit with either mono- or biexponential functions. The fit curves were subsequently differentiated, giving rise to smooth release rate curves from which peak release rates were obtained (**Fig. 2C**). The peak release rates were used to quantify the Ca^2+^ sensitivity of release by dose-response curves.

**Fig. 2.**
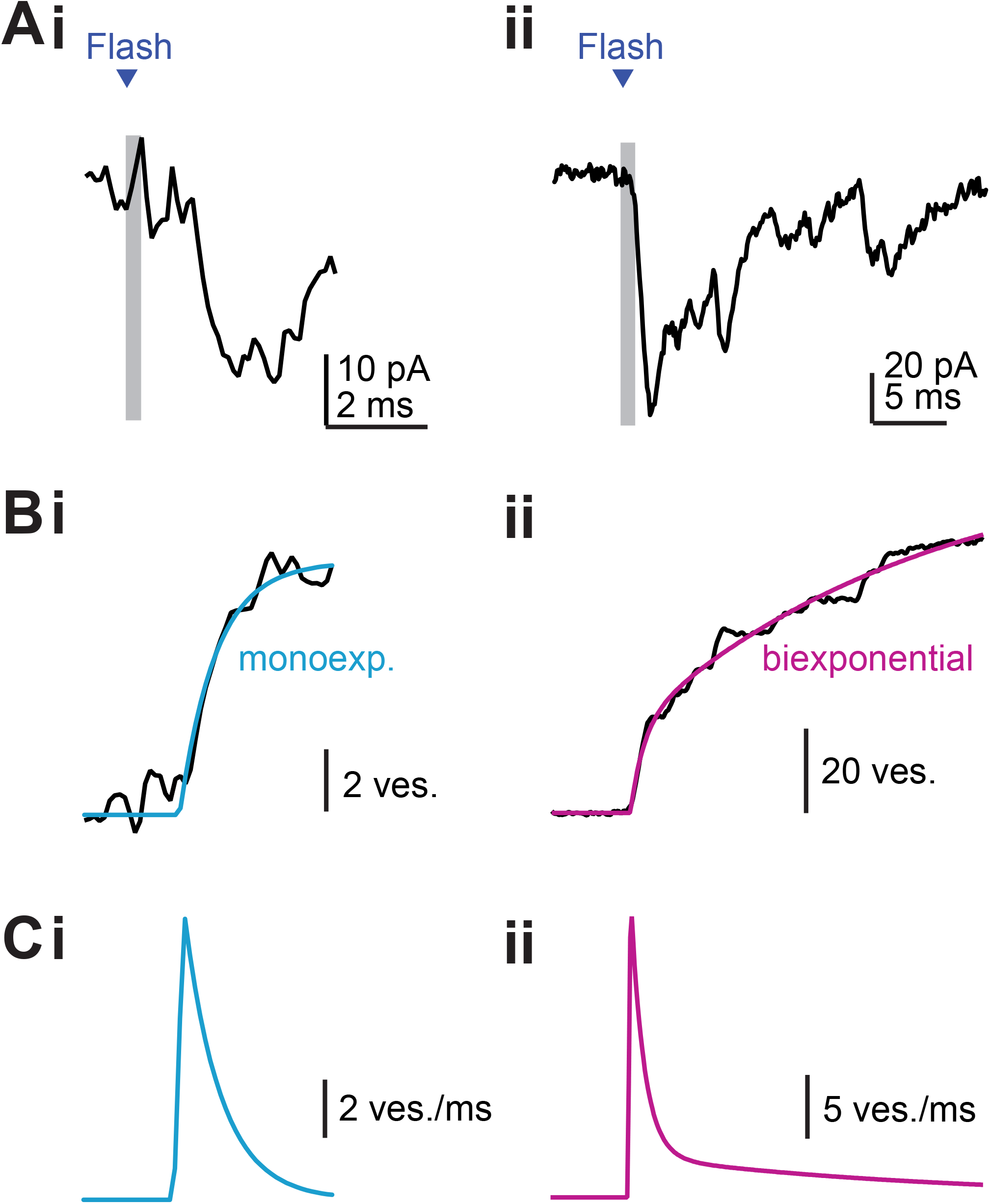
Quantifying release rates from fEPSCs. (**Ai**,**ii**) Examples of fEPSCs evoked by Ca^2+^ uncaging in a connected bouton. UV flashes (grey bars) of 0.2 ms (i) or 1 ms (ii) duration were applied to the bouton. Note the increased fEPSC amplitude and the decreased synaptic delay with increasing flash duration. (**Bi**,**ii**) Release rates were obtained by deconvolution of the fEPSCs with subsequent integration for noise reduction (see **Methods**). The resulting cumulative release rates were fitted either with mono- (i; blue) or biexponential (ii; magenta) functions. (**Ci**,**ii**) The fits from (B) were differentiated resulting in smoothed release rate curves.

Increases in the amplitude and/or duration of the UV-flashes induced increasing Δ[Ca^2+^]_i_ and concomitantly increasing release rates (**Fig. 3A**). fEPSCs were typically detected at presynaptic Δ[Ca^2+^]_i_ above ∼4 μM. Below this value release failures and qEPSCs dominated. Peak release rates increased until Δ[Ca^2+^]_i_ of ∼35 μM with subsequent saturation. The increase was steepest in a narrow dynamic range between ∼10 and ∼30 μM of Δ[Ca^2+^]_i_. Overall, the peak release rate versus Δ[Ca^2+^]_i_ dose-response curve was sigmoidal with positive cooperativity. It was described by a Hill-function with n_H_ of 3.57 and EC_50_ of 20 μM (**Fig. 3B**). While peak release rates increased with Δ[Ca^2+^]_i_, the corresponding synaptic delays decreased (**Fig. 3C**).

**Fig. 3.**
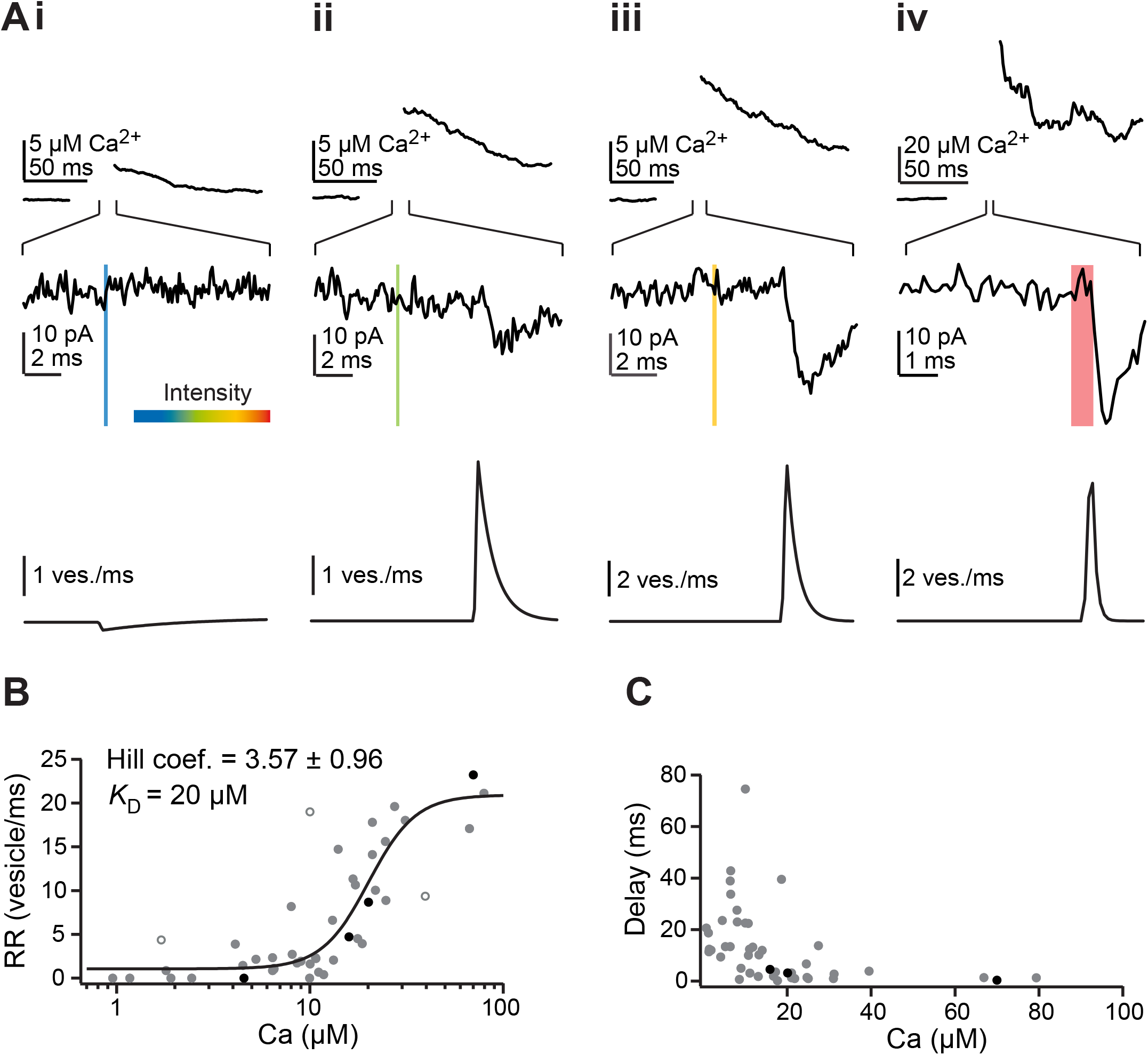
Ca^2+^ dependency of transmitter release from L5PN presynaptic boutons. (**Ai-iv**) UV flashes (colored bars) of different intensities (color code, 0-100%) and durations (bar width) were applied to presynaptic boutons and the corresponding presynaptic Δ[Ca^2+^]_i_ signals (*top*) and release rates (*bottom*) were quantified from simultaneous recordings of the fluorescence signals and the fEPSCs (*middle*) as described in **Figs. 1, 2**. (**B**) Dose-response curve for the dependency of peak release rates (RR) on the corresponding elevations in presynaptic [Ca^2+^]_i_ (n=45, 8 cells). The solid line shows a fit with a Hill function to the data. Open circles indicate outliers that were excluded during fitting. The black points are from the example shown in (A). (**C**) Dependency of synaptic delays on presynaptic Δ[Ca^2+^]_i_.

We tested whether saturation of the peak release rates could result from systematically slower rise times of larger EPSCs (*28*) by plotting peak fEPSC amplitudes against the corresponding peak release rates (**Supplementary Fig. 4**). If larger fEPSCs would have slower rise times, one would expect a shallower relationship between their amplitudes and the corresponding release rates than for smaller fEPSCs. However, we found a linear relationship over the whole range of fEPSCs recorded, indicating that no systematic error is responsible for the saturation of the dose-response curve. Thus, these results indicated substantial differences in the Ca^2+^ -dependency between Syt1- and Syt2-triggered release (see below).

### Photolysis-versus AP-evoked release

To understand whether the dynamic range of the dose-response curve is the physiologically relevant range, we quantified the Ca^2+^-dependency of AP-mediated release (**Fig. 4, Supplementary Fig. 3**). First, we analyzed the dependency of release on the extracellular Ca^2+^ concentration by recording AP-mediated EPSCs in different [Ca^2+^]_e_. The corresponding cumulative release rate curves were typically monoexponential for EPSCs recorded in lower [Ca^2+^]_e_ (≤2mM) and typically became biexponential in higher [Ca^2+^]_e_ (≥5mM). This is reminiscent of the fEPSCs, where lower or higher Δ[Ca^2+^]_i_ resulted in mono- or biexponential cumulative curves, respectively (see above). We plotted the peak amplitudes of the EPSCs and the corresponding peak release rates against the respective [Ca^2+^]_e_. Both curves were sigmoidal, with Hill coefficients similar to those for the intracellular Ca^2+^-dependency of flash-evoked release (**Supplementary Fig. 3Aiii, Biii**). Similar results and parameters for the dependency of AP-evoked release on [Ca^2+^]_e_ were obtained in a previous study on cultured cortical neurons from Syt1 KO mice that were rescued by Syt1 (*15*).

**Fig. 4.**
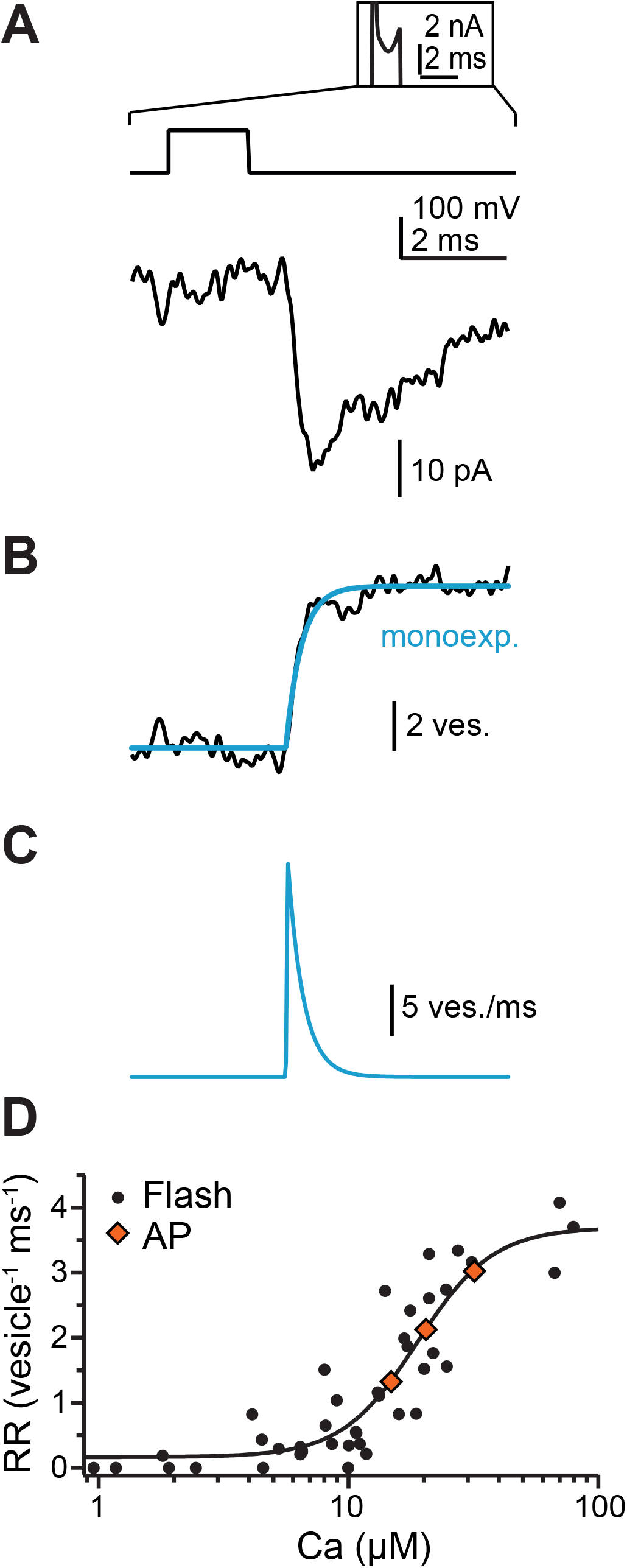
AP evoked release rates. (**A**) Example of an action current (inset, *top*) evoked EPSC (*bottom*) from the same L5PN pair as in **Fig.2**. (**B**) Cumulative release rate obtained from the EPSC in (A) fit with an exponential function. (**C**) Time course of the release rate derived by differentiation of the fit in (B). (**D**) Dose response curve from **Fig.3B** with peak release rates being normalized to the size of the RRP of each bouton. Outlier were omitted (open circles in **Fig.3B**). Diamonds represent AP-mediated peak release rates (25, 50, 75 quartiles) superimposed on the Hill fit.

We previously found that postsynaptic receptor saturation is not an issue at L5PN synapses even under conditions of very high *p*_v_ (*11*). We recorded AP-mediated EPSCs in different [Ca^2+^]_e_ in the presence of γ-D-glutamylglycine (γDGG), which reliefs glutamate receptor saturation and desensitization (*29*). The resulting dose-response curves for EPSC amplitudes and peak release rates were characterized by Hill parameters similar to those obtained in the absence of γDGG (**Supplementary Fig. 3Aiii, Biii**). This suggests, that neither the dose-response curve for the photolysis-evoked release (see above) nor that for the AP-mediated release were strongly affected by postsynaptic receptor saturation.

For more direct comparison between flash- and AP-evoked release we first corrected the photolysis dose-response curve (**Fig. 3B**) for differences in the readily releasable pool of vesicles (RRP) by dividing the peak release rates by the corresponding size of the RRP of each bouton, giving rise to peak release rates per vesicle per ms (*3, 6*) (**Fig. 4D**). The size of the RRP was estimated from the amplitude of cumulative release curves or, in the case of the biexponential curves, from the amplitude of the fast component (*6, 25*). On average, the RRP estimate was 4.9 ± 2.5 (mean ± SD) vesicles. Given that flash-evoked release resulted from the activation of individual boutons (see above) and that the total synaptic connection is formed by 5.5 boutons on average (*26*) we obtain a total RRP of ∼27 vesicles. This is very close to a previous estimate of 28 vesicles obtained for this synapse from cumulative analysis of EPSCs evoked by trains of APs (*30*).

Next, we quantified AP-mediated release rates from a subset of pairs of L5PNs from which the flash-evoked release rates were obtained. We found a median peak release rate of 2.1 vesicle^-1^ ms^-1^ with an IQR of 1.3 – 3.0 vesicle^-1^ ms^-1^ (n=5 cells). Superimposing these values with the Hill fit to the dose-response curve of the flash-induced release rates showed that the local Δ[Ca^2+^]_i_ at Syt1 during an AP reached a median peak value of 20.4 μM and that AP-mediated Δ[Ca^2+^]_i_ span a range of 14.7 - 31.8 μM. (**Fig. 4D**). These data indicate that AP-evoked release rates span the dynamic range of the dose-response curve for photolysis-evoked release with a median value corresponding to its EC_50_. Thus, the dynamic range of the curve indeed reflects the physiologically most relevant range.

### Functional differences between Syt1- and Syt2-triggered release

To gain quantitative insights into the kinetics of Ca^2+^-triggered transmitter release at L5PN synapses and for comparison with Syt2-triggered release, we aimed to establish a kinetic model that describes the experimental photolysis dose-response curve (**Fig. 5**). We first modelled the expected Δ[Ca^2+^]_i_ time course after flash-photolysis, using a detailed model of DMn uncaging (*31*) that included the indicator dye (**Supplementary Fig. 5A**). This model indicated that, similar to previous reports (*2, 25*), under our recording conditions the simulated time course of Δ[Ca^2+^]_i_ was reliably reported by the indicator dye ∼0.5 ms after the flash (**Supplementary Fig. 5B**). Hence, as for the experimental data, for modelling the Ca^2+^-dependency of release we used the Δ[Ca^2+^]_i_ reported by the dye.

**Fig. 5.**
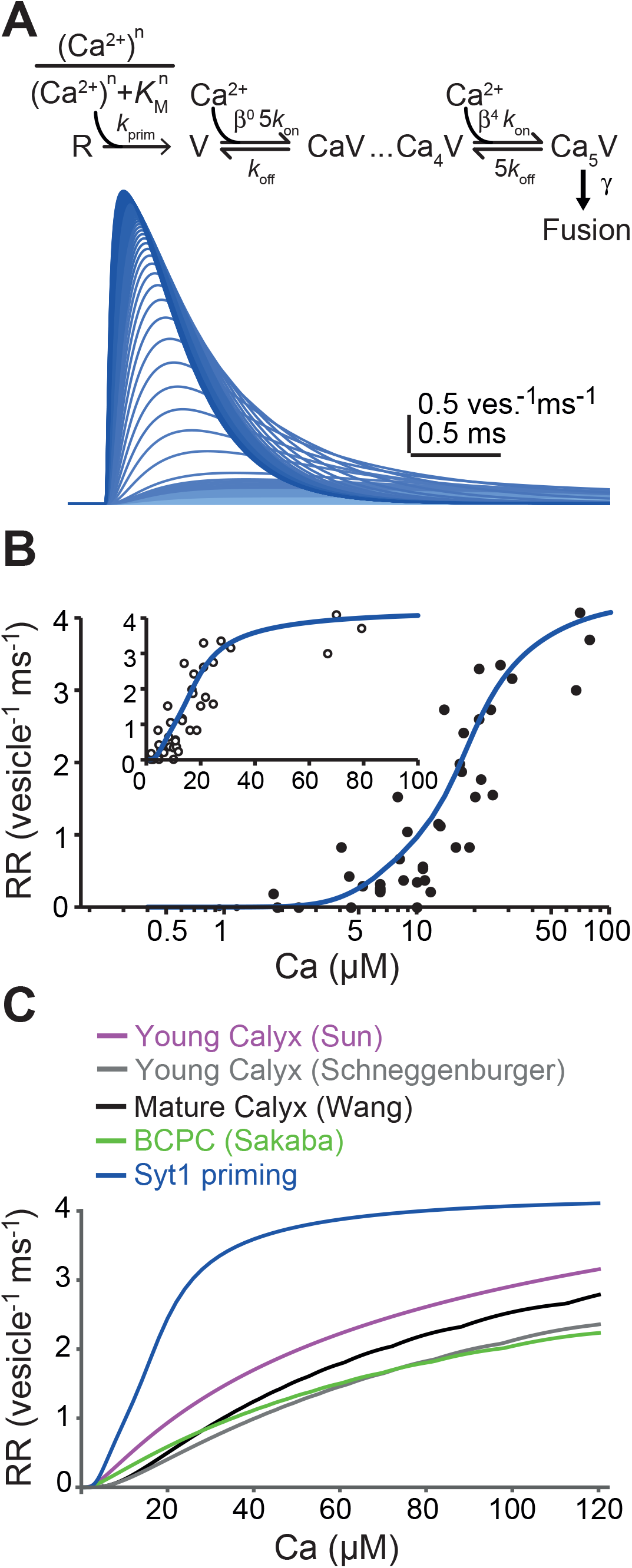
Model of the Ca^2+^-dependency of Syt1-triggered release. (**A**) *Top*: Scheme of the priming model of the Ca^2+^-dependency of release from L5PN boutons. The model covers a Ca^2+^-dependent priming step, five steps for binding of Ca^2+^ to Syt1 with positive cooperativity (Δ) and Ca^2+^-independent fusion from the fully Ca^2+^-occupied sensor (γ). *Bottom*: Release rates simulated for increasing [Ca^2+^]_i_ with the model. (**B**) Fit of the model (blue curve) to the dose-response curve of the peak release rates (RR) from **Fig. 4B**. (**C**) Comparison of the Ca^2+^-dependency of release predicted by the Syt1 priming model (blue) to those of four Syt2 models: Young calyx of Held in gray (*2*) and magenta (*18*), mature calyx (black) (*21*), GABA-ergic basket cell (BC) to Purkinje cell (PC; green) (*6*).

We proceeded by modeling Ca^2+^ binding to Syt1 and release. Like Syt2, Syt1 comprises five Ca^2+^ binding sites (*32*), and the Hill coefficient we obtained is consistent with ≥4 binding steps. Hence, we assumed five Ca^2+^ binding steps for Syt1, followed by fusion from the fully Ca^2+^-bound sensor (**Fig. 5A, Supplementary Fig. 6**). As a starting point we tested previously published Syt2 models (*2, 6, 18, 22*). We found that none of the models could describe our data (**Fig. 5B, C**). All attempts to adjust the parameters to obtain a satisfactory fit to our data failed for any of the Syt2 models. We identified the realization of cooperativity in the Ca^2+^-unbinding rates in the Syt2 models as the main reason. Instead, when we implemented positive cooperativity in the forward binding rates of a five-site model, we readily obtained a model that yielded a much better fit to the dose-response curve as well as to the Ca^2+^-dependency of the synaptic delays (**Supplementary Fig. 6**). The fit to the later phase of the dynamic range and the subsequent saturation could be improved further, if a Ca^2+^-dependent priming step with Michaelis-Menten kinetics was added to the model (**Fig. 5**; Pearsons χ^2^ test; Δpχ^2^=0.27 between the two models). This may indicate that during larger Ca^2+^-elevations (> ∼18-20 μM) release is boosted via vesicle recruitment, which is consistent with the occasional occurrence of biexponential fEPSCs at larger Ca^2+^ elevations (see above). It appears that both processes, release and recruitment, cannot be accelerated further by Ca^2+^ elevations above ∼35 μM.

The Ca^2+^ dependencies of release as reported by the Syt2 models were shifted towards lower affinity compared to our Syt1 data and model. Hill fits to the simulations with the Syt2 models shown in **Fig. 5C** gave EC_50_ values between 63 and 82 μM. In addition, the Ca^2+^-dependency of Syt2-triggered release typically gets shallower above ∼5 to 10 μM Δ[Ca^2+^]_i_ (*2, 3, 21*) and, hence, the Syt2 models did not fit the dynamic range of Syt1-triggered release (**Fig. 5C**). These results indicate that Syt1-triggered release from L5PN boutons has three to four-fold higher Ca^2+^ sensitivity with a stronger Ca^2+^-dependency in the dynamic range than Syt2-triggered release.

## Discussion

To our knowledge our data provide the first kinetic description of the Ca^2+^-dependency of Syt1-triggered release from synaptic boutons in intact brain tissue. They explain major differences in release between neocortical and hindbrain synapses, in particular the high *p*_v_ (∼0.63) and high synaptic reliability observed in the neocortex (*11, 26, 33-35*). Furthermore, the dynamic range could provide a framework for enhanced plasticity induced by Ca^2+^ buffers (**Supplementary Fig. 7, 8**; see below).

The *K*_D_ of 20 μM we found is ∼10-fold lower than for the isolated C2B domain of Syt1, but ∼4-fold higher than for the C2B domain in the presence of PIP_2_ (*36*). Hence, the details of the context into which Syt1 is embedded are significant for determining the kinetics of release. *In vivo*, Ca^2+^ sensors are integrated in a supra-molecular protein complex and interact with lipids, which further influences their Ca^2+^-binding kinetics (*32, 36-38*). Thus, it was required to quantify the Ca^2+^ sensitivity of Syt1-triggered release at synaptic boutons in brain tissue with intact presynaptic release machinery. Indeed, our results show that the rate constants for Ca^2+^-binding and fusion in neocortical boutons are orders of magnitude higher than those of Syt1-triggered fusion of dense core vesicles in chromaffin cells (*39, 40*).

The analysis of the Ca^2+^-dependency of release has previously been restricted to synapses expressing Syt2 (*2, 3, 6, 18, 19, 25*). Our results show that the Ca^2+^-dependency of Syt1-triggered release deviates from that reported for Syt2. First, Syt1-triggered release had at least three-fold higher affinity. Second, the dose-response curve for Syt1 was steeper in the dynamic range between ∼10 and ∼30 μM Δ[Ca^2+^]_i_ than the corresponding Syt2 curves (**Fig. 5C**). Finally, Syt1-triggered release showed pronounced saturation above the dynamic range, while the Syt2 simulations show much slower saturation. Overall, these differences necessitated establishing a new model to describe the Ca^2+^-dependency of Syt1-triggered release (**Fig. 5A**).

The estimates of peak release rates and the degree of saturation can be influenced by priming (*6, 25*). At synapses with rapid vesicle recruitment and sizable replenishment pools this can even prevent dose-response curves from saturation (*25*). We found that also the Syt1 dose-response curve was better described in the upper range of the Δ[Ca^2+^]_i_ by a model that incorporated a Ca^2+^-dependent priming step (**Fig. 5, Supplementary Fig. 6**). Yet, priming did not prevent pronounced saturation of the curve. This is consistent with observations suggesting that the immediate replenishment pool at mature L5PN boutons is rather limited (*30*).

Release sensor models are important tools for quantifying functional presynaptic nanotopographies (*4, 5, 8-13*). We previously estimated the nanodomain coupling distance between a single Ca_v_2.1 channel and the vesicular release sensor to be ∼7 nm (*11*) at the L5PN synapse using a Syt2 model (*22*). With the new Syt1 models we refine this estimate to a range of 11-16 nm (**Supplementary Fig. 7**), which is still a tight nanodomain coupling. The corresponding estimates for the amplitudes of the local AP-mediated Δ[Ca^2+^]_i_ at the sensor are 35-21 μM. In line with this estimate of Δ[Ca^2+^]_i_, we found here that AP-mediated release rates cover the dynamic range of the Syt1 dose-response curve with a median value almost identical to the EC_50_ of 20 μM. Thus, AP-mediated release from neocortical boutons will be highly sensitive to modulations in Δ[Ca^2+^]_i_ in the dynamic range.

To further pursue this notion, we incorporated either Syt1 or Syt2 into previously published models of L5PN active zones (*11*) with nanodomain or microdomain coupling (**Supplementary Fig. 8**). Irrespective of the coupling topography, Syt1-triggered release was less sensitive to variations in the channel to sensor distances than Syt2-triggered release. This is due to the higher Ca^2+^-affinity of Syt1 and provides an explanation for the high reliability of neocortical synapses that was found to be independent of age and the details of the presynaptic coupling topography (*11, 33, 41*). In loose microdomain coupling, which was found in the young developing neocortex (*11, 41*), Syt1-triggered release was ∼2 fold more sensitive to the presence of a slow Ca^2+^ buffer than Syt2-triggered release (**Supplementary Fig. 8B**). This is due to the steep slope of the Ca^2+^ sensitivity of Syt1 in the dynamic range. For Syt2-triggered release, loose microdomain coupling was previously shown to provide a framework for enhanced plasticity resulting from Ca^2+^ buffering (*13*). We show here that the properties of Syt1 further expand the possibilities for plastic regulation of release via Ca^2+^ buffering in the regime of loose microdomain coupling. Thus, the combination of high affinity with steep Ca^2+^-dependency of Syt1-triggered release can promote both, high-fidelity synaptic transmission at moderate local presynaptic Ca^2+^ elevations as well as enhanced plasticity. This will make neocortical synapses more complex computational devices, which could expand the computational power of cortical microcircuits.

## Acknowledgments

We thank Gudrun Bethge for technical assistance.

## Funding

This work was supported by a grant of the German research foundation (DFG) to HS (SCHM1838/2-1).

## Author contributions

Conceptualization: HS; Funding acquisition: HS, JE; Methodology: all authors; Investigation: GB, SB, AE; Data Analysis: GB, SB, HS; Resources: JE, RK, HS; Modeling: SH, HS; Supervision: HS; Writing – original draft: GB, SB, HS; Writing – review & editing: all authors.

## Competing interests

Authors declare that they have no competing interests.

## Supplementary Materials

### Materials and Methods

#### Slice preparation, electrophysiological paired recordings, and deconvolution of EPSCs

As described previously(*11*) C57BL/6J mice at P21-24 of either sex were decapitated under deep Isoflurane (Curamed) inhalation anesthesia. The brain was excised rapidly and placed in cooled (0-4°C) artificial cerebrospinal fluid (ACSF) containing (in mM): 125 NaCl, 2.5 KCl, 1.25 NaH_2_PO_4_, 26 NaHCO_3_, 1 MgCl_2_, 2 CaCl_2_, and 20 glucose, equilibrated with 95% O_2_ and 5% CO_2_ (pH 7.3-7.4). Neocortical slices (coronar, 250-300 μm thick) were cut from the S1 region with a vibratome (HM 650 V, Microm), incubated for 30 min at 35°C and subsequently stored at room temperature (21°C). For experiments slices were transferred to a recording chamber and continuously perfused with ACSF (2-3 ml per min; supplemented with 10 mM (-)-bicuculline methiodide (Tocris)) at 30-32°C. Unless stated otherwise, chemicals were from Sigma-Aldrich.

Patch pipettes were prepared from borosilicate glass (Hilgenberg) with a PC-10 puller (Narishige) and had final resistances of 6-8 MΩ when filled with the following standard pipette solution (in mM): 150 K-gluconate, 4 NaCl, 3 MgCl_2_, 3 Na_2_ATP, 0.3 NaGTP, 0.05 EGTA, 10 KHEPES, dissolved in purified water. The pH was adjusted to 7.3 with KOH.

Patch-clamp recordings from pairs of L5PNs were performed under optical control (BX51WI, Olympus), using an EPC10/2 amplifier and Patchmaster software (version v2×3.2, HEKA). EPSCs were recorded in the whole-cell configuration at a holding potential (V_hold_) of -80 mV (online corrected for a liquid junction potential of 16 mV), filtered at 5 kHz and sampled at 10 kHz. Series resistance (R_s_) was continuously compensated to a fixed value between 10 and 15 MΩ. Holding current (I_hold_) was monitored continuously. Experiments were rejected when R_s_ or I_leak_ exceeded 30 MΩ or 500 pA, respectively.

Transmitter release rates were derived from the multiquantal EPSCs by the deconvolution method (*2, 27*), using AP-evoked qEPSCs recorded in 0.5 and 1 mM [Ca^2+^]_e_, respectively, as described in detail previously (*11*). Since qEPSCs decayed monoexponentially, the transmitter release rates were calculated from the EPSC, its time derivative and the average amplitude (6 pA) and decay time constant (2 ms) of the qEPSC (**Supplementary Fig. 3A**) according to the equation given in ref. 22. The deconvolution method assumes linear summation of the qEPSCs (*2, 27*). Due to dendritic filtering the values of the peak release rates likely represent lower limit estimates (*2*).

#### Ca^2+^ uncaging and Ca^2+^ imaging

For Ca^2+^ uncaging experiments presynaptic L5PNs were loaded with an uncaging pipette solution containing (in mM): 10 DMn, 10 x F CaCl_2_, 0.8 OGB5N, 0.1 Atto594, 130 KCl, 6 NaCl, 0.5 MgCl_2_, 3 Na_2_ATP, 0.3 NaGTP, 20 KHEPES, dissolved in purified water. The pH was adjusted to 7.3 with KOH and DMn was saturated with equimolar concentrations of CaCl_2_ depending on DMn purity (F). The purity of each DMn batch was determined in purified water through sequential addition of Ca^2+^ as previously described (*42*) and by measuring the free Ca^2+^ concentration using the dual dye method (10 μM OGB1, 5 μM Atto594). Fractional purity of DMn was typically 0.7 to 0.8. Pipette solutions for minimal (R_min_) or maximal G/R ratios (R_max_) were prepared by replacing CaCl_2_ in the uncaging solution by 10 mM K_2_EGTA or by increasing the CaCl_2_ concentration to 25 mM, respectively. In order to reduce spontaneous activity and a flash-artifact in the electrophysiological recordings (*25*), prior to uncaging 1 μM TTX and 20 mM TEA were added to the bathing solution in most experiments.

Presynaptic fluorescence signals were elicited by UV-flashes (0.1 to 1 ms duration, 25 to 100% intensity) and recorded with a two-photon microscope in the point mode at 500 kHz temporal resolution with subsequent binning to 500 Hz (**Fig. 1A**). For probing signal homogeneity among boutons, line scans at 500 Hz resolution were used (**Supplementary Fig. 2**). A custom-build two-photon microscope based on a Fluoview-300 scanner (Olympus), a 60x/0.9 N.A. objective, a mode-locked Ti:sapphire laser (Tsunami, Newport-Spectra Physics, set to a center wavelength of 810 nm), a Pockels cell (350-80 KD*P, Conoptics) and a DL 375/200 UV laser system (375 nm, 200 mW, Rapp Optoelectronic) was used. During UV-flashes the photomultiplier tubes (PMTs) detecting the fluorescence-photons were protected by a shutter (**Fig. 1A**).

The profile of the UV laser illumination at the specimen plane was assessed *in vitro* by uncaging of fluorescein (CMNB-caged fluorescein, Thermo Fisher Scientific) as described previously (*25*). To limit the mobility of the UV-released fluorescein, Agar blocks (1.5%) were prepared with 100 μM caged fluorescein(*3, 25*). The x-y fluorescence profile of the dye after being released from the cage was measured at different z-positions. The intensity of fluorescein was homogenous within a volume of 10 μm * 10 μm * 5 μm (x*y*z), which fully encompasses the small L5PN boutons (radius ∼0.25 to 0.5 μm(*26, 43, 44*)). Outside this region, the intensity rapidly dropped over a distance of ∼5 μm to baseline fluorescence. Care was taken that in z-direction no other boutons except for those in the focal plane could have contributed to the fEPSCs. The focal volume of our two-photon system was quantified with fluorescent nanospheres (radius xy = 0.53 μm; radius z = 1.95 μm (*45*)) and is equal to or even larger than the size of L5PN terminals. Hence, the fluorescence represents volume averaged signals and within the bouton volume the Ca^2+^ elevations can be expected to be homogeneous.

The fluorescence signals were filtered (HC647/75, Semrock HC525/50, 720-SP, AHF), detected by two external PMT modules (H7422-40, Hamamatsu; PMT-02M/PMM-03, NPI electronics) monitoring red and green epi-fluorescence, respectively at fixed PMT voltages, and digitized with the Fluoview system. The Ca^2+^-dependent green fluorescence was normalized to the Ca^2+^-insensitive red fluorescence and expressed as background-corrected ΔG/R (*46*). The Background subtraction removed a residual flash-artifact in the fluorescence recordings (**Fig. 1**) (*25*). To account for a slight continuous bleaching in the red fluorescence that was not evident in the green fluorescence, we used a linear correction algorithm. ΔG/R signals were converted to Δ[Ca^2+^]_i_ based on an *in vitro* determination of *K*_D_ values and the determination of R_min_ and R_max_ values (**Supplementary Fig. 1**) (*47*). The *K*_D_ value of the Ca^2+^ indicator dye was quantified from *in vitro* calibrations in a commercial buffer kit (Biotium) and in our pipette solution adjusted with CaEGTA and K_2_EGTA (100 mM stock solutions, 10 mM HEPES added) to contain different free Ca^2+^ concentrations that were calculated with the MaxChelator(https://somapp.ucdmc.ucdavis.edu/pharmacology/bers/maxchelator/CaMgATPEGTA-NIST.htm). Calibration in the buffer kit and in KCl-based pipette solution yielded similar *K*_D_ values for OGB5N (*K*_D_ = 29.8 μM and 22.0 μM respectively; **Supplementary Fig. 1A,B**). R_min_ and R_max_ values were measured after each successful paired recording in sealed patch pipettes that were placed in the slices near the recorded cells. The pipettes contained the KCl-based uncaging solution with either 0 Ca^2+^ and 10 mM EGTA (zero Ca^2+^ solution) or 25 mM free Ca^2+^ (high Ca^2+^ solution). Since DMn contributes to the fluorescence, R_min_ and R_max_, were determined in the presence of the experimental DMn concentrations. In addition, the fluorescence properties of DMn change after photolysis, and the Ca^2+^ sensitive and insensitive dyes can differentially bleach during UV and two-photon illumination (*42, 48, 49*). Hence, R_min_ and R_max_ values were determined before and after a flash, using the same flash intensities and durations as during the experiments. While R_max_ values were affected by the flash, the R_min_ values remained stable (*25, 42, 49*) (**Supplementary Fig. 1C**). We assumed no effect of the UV flash on the *K*_D_ of the Ca^2+^ indicator dyes (*50*).

### Data analysis and statistics

Electrophysiological and Ca^2+^ imaging data were analyzed with custom written routines in Igor Pro 8 (Wavemetrics). Data are expressed as median and interquartile range (IQR), except for dye calibrations, which are shown as mean ± standard deviation (SD). In boxplots mean values are included additionally as dashed lines. Since samples sizes were small, all data were compared with the more restrictive non-parametric tests, using the Mann-Whitney-U rank sum test (MWU; two groups) or an ANOVA on ranks (more than two groups). Paired data were compared with the Wilcoxon signed rank test. All statistical tests were two-tailed. P values are indicated as *P ≤ 0.05, **P ≤ 0.01 and ***P ≤ 0.001. The number of experiments (n) represents the number of cells or boutons as indicated and was chosen to be sufficiently high for concise statistical analysis. Statistics were performed with Sigma Plot 11.0 (Dundas Software), Graphpad Prism 10 and Mathematica 11.

### Modeling

For modeling Ca^2+^ dynamics and release, the kinetic reaction schemes for Ca^2+^ and Mg^2+^ uncaging and -binding (**Fig. 5, Supplementary Figs. 5, 7**) were converted to systems of ordinary differential equations (ODEs) that were numerically solved using “NDSolve” or “NDSolveValue” in Mathematica 13 (Wolfram) as described previously (*11, 25*). For modeling release the initial conditions were calculated for the resting [Ca^2+^]_i_ similar to the uncaging simulation (see below) and subsequently the model was driven by the Δ[Ca^2+^]_i_. For modeling the Ca^2+^ dynamics within a bouton, a single compartment model was used due to the expected homogenous Ca^2+^ elevations in the boutons. The initial conditions for the uncaging simulation were derived as follows: First, the system of ODEs was solved for the steady-state, using total concentrations of all species and the experimentally determined values of the pre-flash resting [Ca^2+^]_i_ as starting values. Subsequently, the values obtained for all free and bound species were used as initial conditions for the uncaging simulation. The kinetic properties of DMn were simulated according to Faas et al. (*51, 52*). The total DMn concentration ([DMn]_T_) consists of the free form ([DMn]), the Ca^2+^ bound form ([CaDMn]), and the Mg^2+^ bound form ([MgDMn]). Each of these forms subdivided into an uncaging fraction (α) and a non-uncaging fraction (1-α). The uncaging fraction further subdivided into a fast (af) and a slow (1-af) uncaging fraction. This is described by the following equations:

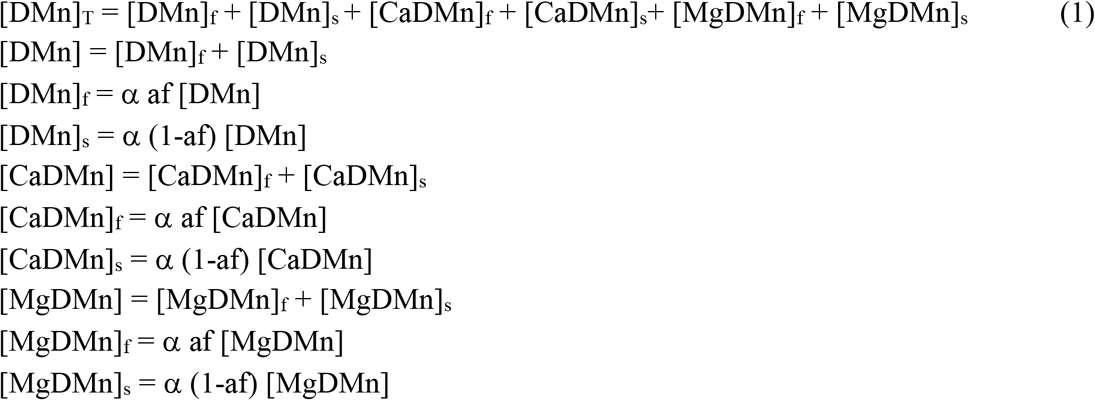

The suffixes “T”, “f”, and “s” stand for “total”, “fast” and “slow”, respectively. The transition of fast and slow uncaging fractions into low affinity photoproducts (PP) occurred with fast (τ_f_) or slow (τ_s_) time constants, respectively. Free, Ca^2+^ or Mg^2+^ -bound DMn decomposed into two PP (PP1, PP2) with different binding kinetics. The metal-binding kinetics of all species were governed by the corresponding forward (*k*_on_) and backward (*k*_off_) rate constants

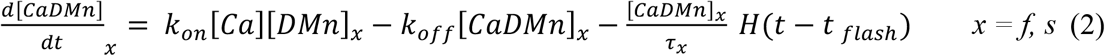

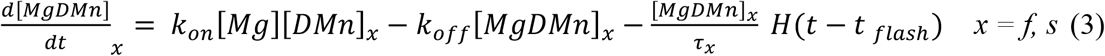

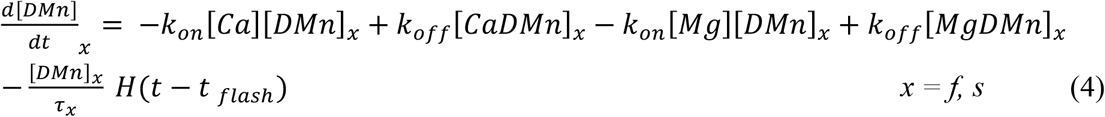

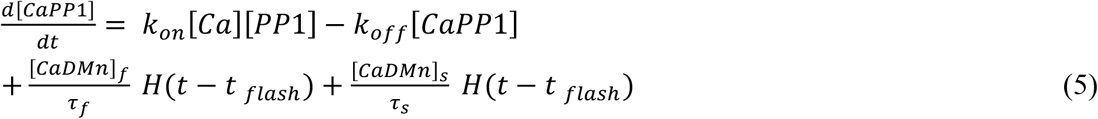

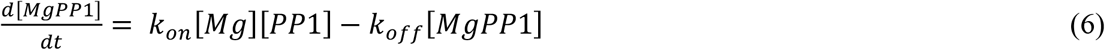

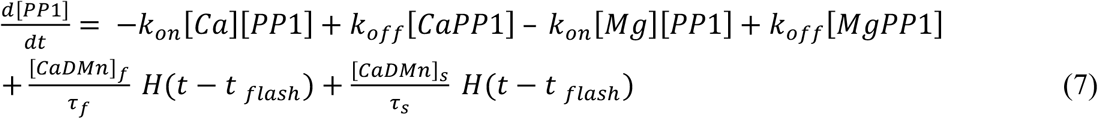

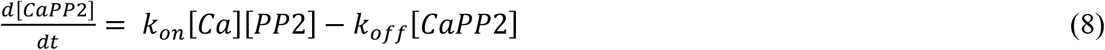

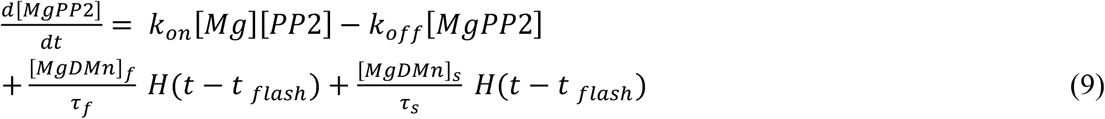

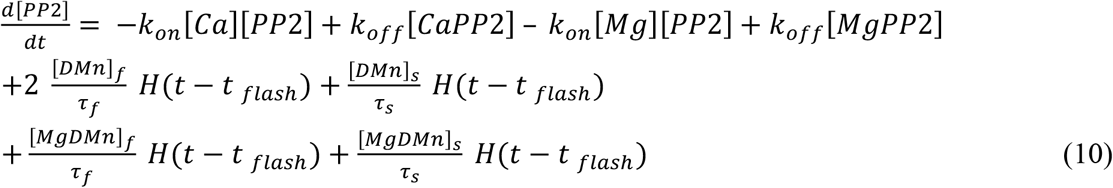

where H is the Heaviside step function and t_flash_ the time of the UV flash. Ca^2+^ and Mg^2+^ binding to the dye, ATP and an endogenous buffer (EB) were simulated by second order kinetics for all buffers (B):

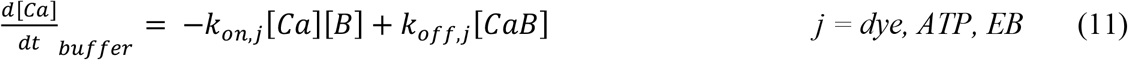

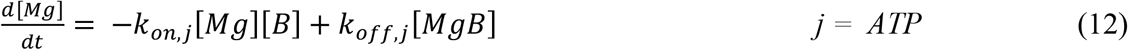

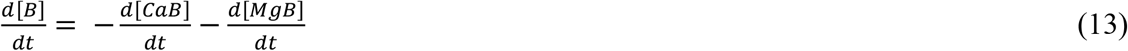

For clearing of Ca^2+^ from the cytosol, a linear pump with rate γ was simulated:

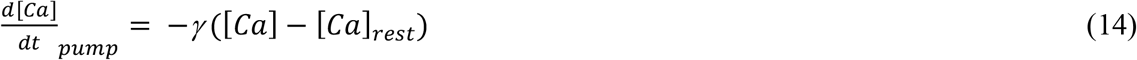

The time course of the total change in [Ca^2+^]_i_ or [Mg^2+^]_i_ is given by the sum of all above equations involving changes in [Ca^2+^]_i_ or [Mg^2+^]_i_, respectively. [Ca^2+^]_i_ as reported by the dye was calculated from the concentration of the Ca^2+^-dye complex assuming equilibrium conditions (*53*). For fitting the simulation to the experimental Ca^2+^ transients, α and γ were the free parameters. The parameters of the model are given in the **Supplementary Table 1**.

Under certain conditions, Ca^2+^-uncaging was reported to lead to brief spike-like overshoots in the first millisecond of the Δ[Ca^2+^]_i_ time course (*2, 3, 31*) that would have remained undetected by the Ca^2+^ imaging (see above). Under our recording conditions a predicted initial spike-like overshoot in Δ[Ca^2+^]_i_ was not prominent. After ∼0.5 ms the dye reliably reflected the time course of [Ca^2+^]_i_ (**Supplementary Fig. 5**), which is similar to other previous reports (*2, 25, 31*).

**Supplementary Table 1.** Model Parameters.

| Parameters |  | Values | References/Notes |
| --- | --- | --- | --- |
| Total magnesium | $[\text{Mg}^{2+}]_T$ | $0.5 \cdot 10^{-3} \text{ M}$ | Pipette concentration |
| OGB-5N | [OGB] | $200\text{-}800 \cdot 10^{-6} \text{ M}$ | Pipette concentration |
| | $K_D$ | $22 \cdot 10^{-6} \text{ M}$ | Measured |
| | $k_{\text{off}}$ | $5720 \text{ s}^{-1}$ | Calculated |
| | $k_{\text{on}}$ | $2.6 \cdot 10^8 \text{ M}^{-1} \text{ s}^{-1}$ | (52, 54) |
| ATP | [ATP] | $3 \cdot 10^{-3} \text{ M}$ | Pipette concentration |
| Ca<sup>2+</sup> binding | $K_D$ | $2 \cdot 10^{-4} \text{ M}$ | (55) |
| | $k_{\text{off}}$ | $100\,000 \text{ s}^{-1}$ | ibid. |
| | $k_{\text{on}}$ | $5 \cdot 10^8 \text{ M}^{-1} \text{ s}^{-1}$ | ibid. |
| Mg<sup>2+</sup> binding | $K_D$ | $100 \cdot 10^{-6} \text{ M}$ | (3); MaxC |
| | $k_{\text{off}}$ | $1000 \text{ s}^{-1}$ | ibid. |
| | $k_{\text{on}}$ | $1 \cdot 10^7 \text{ M}^{-1} \text{ s}^{-1}$ | ibid. |
| Endogenous buffer | [EB] | $50 \cdot 10^{-6} \text{ M}$ | (11, 44) |
| | $K_D$ | $32 \cdot 10^{-6} \text{ M}$ | calculated |
| | $k_{\text{off}}$ | $1000 \text{ s}^{-1}$ | (11, 13) |
| | $k_{\text{on}}$ | $5 \cdot 10^8 \text{ M}^{-1} \text{ s}^{-1}$ | ibid. |
| Total DM nitrophen | $[\text{DMn}]_T$ | $10 \cdot 10^{-3} \text{ M}$ | Pipette concentration |
| Ca<sup>2+</sup> binding | $K_D$ | $6.5 \cdot 10^{-9} \text{ M}$ | (51) |
| | $k_{\text{off}}$ | $0.19 \text{ s}^{-1}$ | ibid. |
| | $k_{\text{on}}$ | $2.9 \cdot 10^7 \text{ M}^{-1} \text{ s}^{-1}$ | ibid. |
| Mg<sup>2+</sup> binding | $K_D$ | $1.5 \cdot 10^{-6} \text{ M}$ | ibid. |
| | $k_{\text{off}}$ | $0.2 \text{ s}^{-1}$ | ibid. |
| Uncaging fraction | $\alpha$ | variable | |
| Fast uncaging fraction | af | 0.67 | (51) |
| Photoproduct 1 | [PP1] |  |  |
| Ca<sup>2+</sup> binding | $K_D$ | $2.38 \cdot 10^{-3} \text{ M}$ | (51) |
| | $k_{\text{off}}$ | $69\,000 \text{ s}^{-1}$ | ibid. |
| | $k_{\text{on}}$ | $2.9 \cdot 10^7 \text{ M}^{-1} \text{ s}^{-1}$ | ibid. |
| Mg<sup>2+</sup> binding | $K_D$ | $1.5 \cdot 10^{-6} \text{ M}$ | ibid. |
| | $k_{\text{off}}$ | $300 \text{ s}^{-1}$ | ibid. |
| | $k_{\text{on}}$ | $1.3 \cdot 10^5 \text{ M}^{-1} \text{ s}^{-1}$ | ibid. |
| Photoproduct 2 | [PP2] |  |  |
| Ca<sup>2+</sup> binding | $K_D$ | $124.1 \cdot 10^{-6} \text{ M}$ | ibid. |
| | $k_{off}$ | $3600 \text{ s}^{-1}$ | ibid. |
| | $k_{on}$ | $2.9 \cdot 10^7 \text{ M}^{-1} \text{ s}^{-1}$ | ibid. |
| $\text{Mg}^{2+}$ binding | $K_D$ | $1.5 \cdot 10^{-6} \text{ M}$ | ibid. |
| | $k_{off}$ | $300 \text{ s}^{-1}$ | ibid. |
| | $k_{on}$ | $1.3 \cdot 10^5 \text{ M}^{-1} \text{ s}^{-1}$ | ibid. |
| Release sensor model |  |  |  |
| ‘Five-site’ | $k_{off}$ | $10\,000 \text{ s}^{-1}$ | Fit parameter |
| | $k_{on}$ | $1 \cdot 10^8 \text{ M}^{-1} \text{ s}^{-1}$ | ditto |
| | $\gamma$ | $5900 \text{ s}^{-1}$ | ditto |
| | $\beta$ | 4 | Cooperativity factor |
| ‘Priming’ | $k_{off}$ | $10\,000 \text{ s}^{-1}$ | Fit parameter |
| | $k_{on}$ | $1 \cdot 10^8 \text{ M}^{-1} \text{ s}^{-1}$ | ditto |
| | $\gamma$ | $4000 \text{ s}^{-1}$ | ditto |
| | $\beta$ | 4 | Cooperativity factor |
| | $K_M$ | $20 \cdot 10^{-6} \text{ M}$ | Fit parameter |
| | $k_{prim}$ | $6000 \text{ s}^{-1}$ | ditto |
| | $n$ | 5 | Fit parameter |

## Supplementary Figure Legends

**Supplementary Fig. 1.**
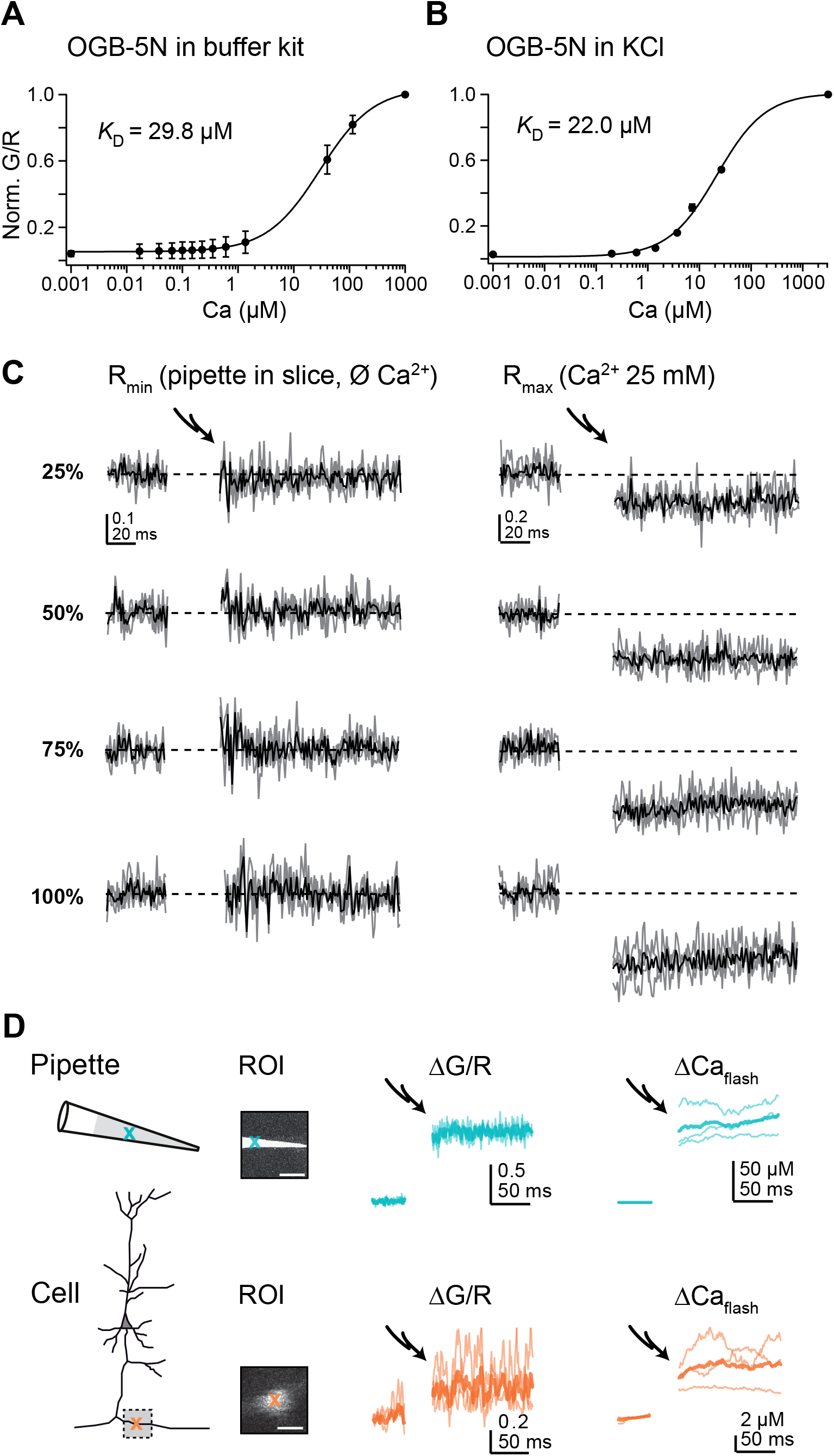
*In vitro* calibration curves and conversion of ΔG/R to Δ[Ca^2+^]_i_. (**A**) Calibration curve of OGB5N in a KCl-based commercial buffer kit. Each data point is an average of 3 measurements (mean ± SD). The *K*_D_ value was derived from a Hill fit (solid curve). (**B**) As in (A) but for our KCl-based pipette solution (n=2). (**C**) UV flashes of different intensities were applied to sealed pipettes containing either R_min_ (0 Ca^2+^) or R_max_ solutions (25 mM Ca^2+^). The pipettes were placed into slices at positions near recorded cells. (**D**) Ca^2+^ uncaging in a pipette and a bouton. *Top*: Scheme of a pipette with indicated region of interest (ROI) from which pre- and post-flash (0.1ms 100%) ΔG/R ratios (middle) were measured in point scans (cross) and converted to Δ[Ca^2+^]_i_ (right) based on the cuvette calibrations. *Bottom*: Scheme of a L5PN with indicated ROI from which ΔG/R were measured (middle) and converted to the corresponding flash-induced Δ[Ca^2+^]_i_ (right).

**Supplementary Fig. 2.**
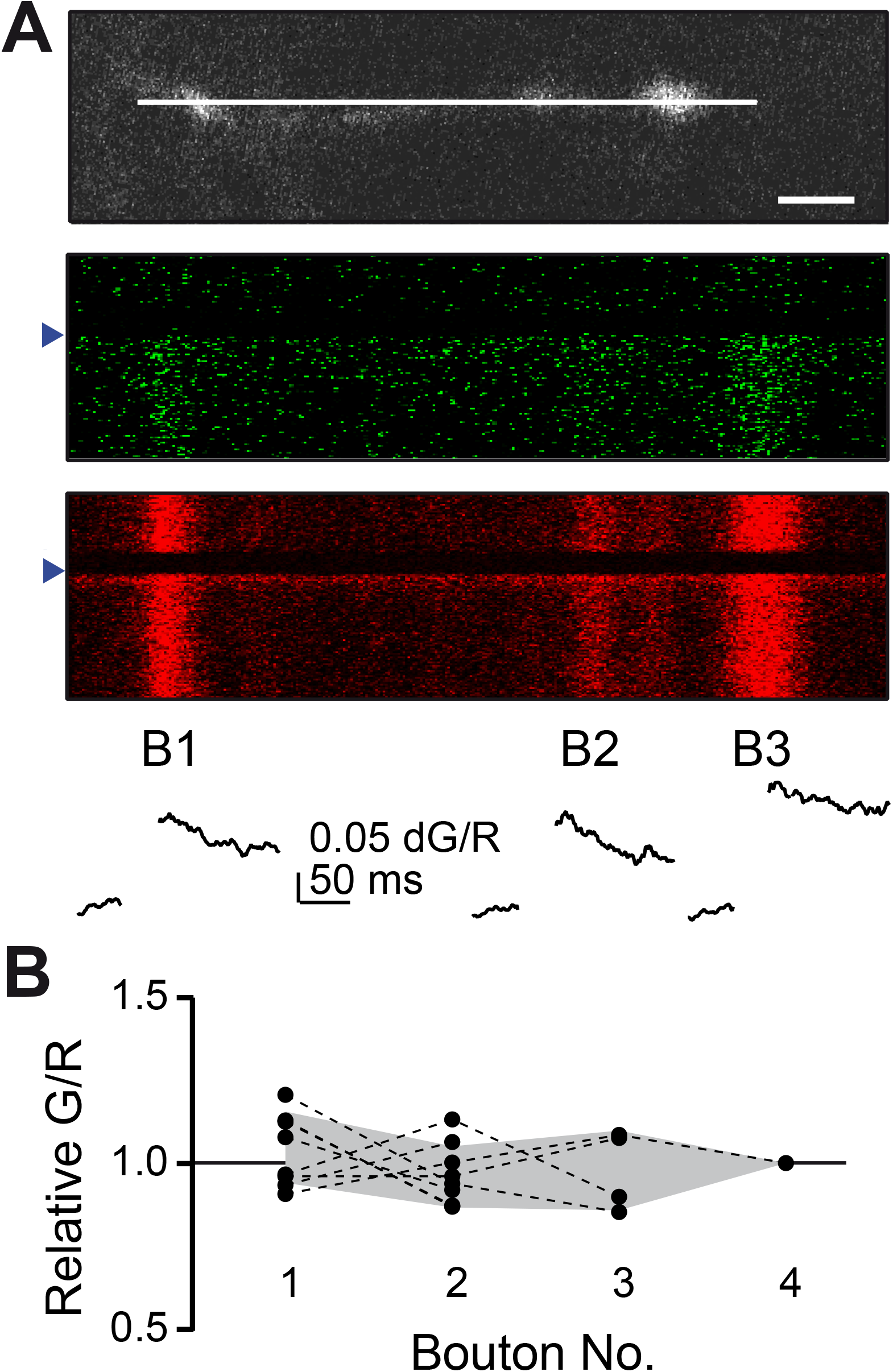
Homogeneity of flash-induced fluorescence signals among boutons. (**A**) UV-flash (triangle) induced G and R signals (*middle, bottom*) were recorded simultaneously in 2-4 boutons in line-scans (*top*, scale bar 2 μm). (**B**) Summary of G/R ratios (normalized to the median G/R of given scan; IQR in grey) from 14 line scans. These recordings indicate homogeneity of flash-induced elevations in [Ca^2+^]_i_ among boutons of a given scan.

**Supplementary Fig. 3.**
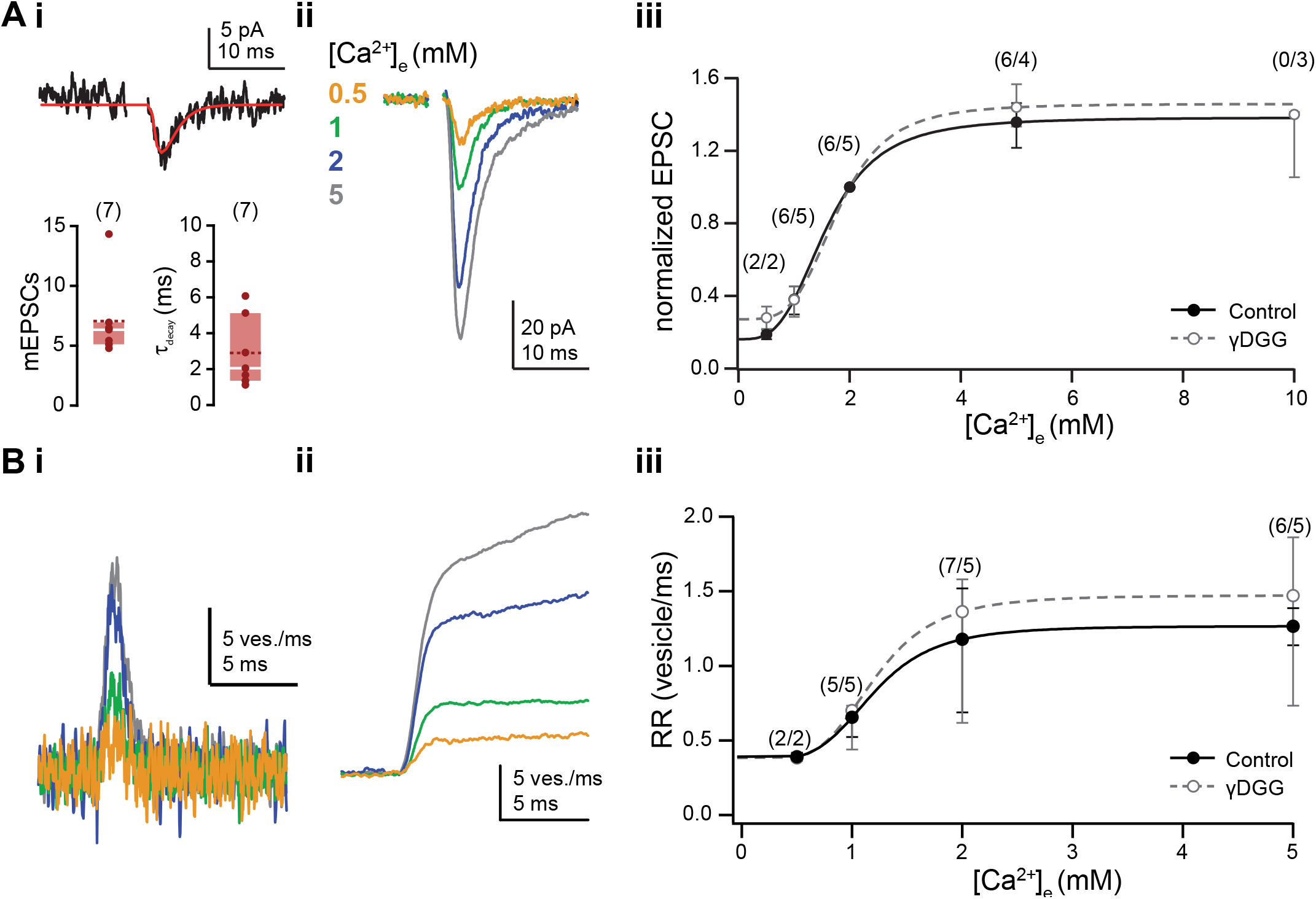
qEPSCs and dose-response curves for AP-mediated release. (**Ai**) *Top*: Example of an evoked qEPSCs recorded from a pair of L5PNs in 0.5 mM [Ca^2+^]_e_ and fitted with an alpha function (red). *Bottom*: Averaged qEPSC amplitudes and decay time constants (τ_decay_) determined from alpha fits. (**Aii**) Examples of averaged EPSCs (n>10 individual recordings each) recorded from a pair of L5PNs (P22) in the indicated [Ca^2+^]_e_. (**Aiii**) EPSC amplitudes were normalized to amplitudes in 2 mM [Ca^2+^]_e_ and plotted against the corresponding [Ca^2+^]_e_ (filled circles; median and IQR). The solid curve represents a fit with a Hill function (*K*_D_, 1.58 mM, Hill coefficient 3.32). To test for postsynaptic receptor saturation and desensitization EPSCs were also recorded in the presence of 1-2 mM γDGG (open circles). The dashed curve is the corresponding Hill fit (*K*_D_, 1.78 mM, Hill coefficient 3.99). (**Bi**,**ii**) Release rates (i) and their temporal integrals (ii) calculated by deconvolution of the EPSCs in (Aii). (**Biii**) Peak release rates (RR) derived by deconvolution of EPSCs recorded in the absence (filled circles) and presence (open circles) of γDGG were plotted against the corresponding [Ca^2+^]_e_ and fitted with Hill functions (solid line: *K*_D_, 1.21 mM, Hill coefficient 4.29; dashed line: with γDGG, *K*_D_, 1.20 mM, Hill coefficient 4.37).

**Supplementary Fig. 4.**
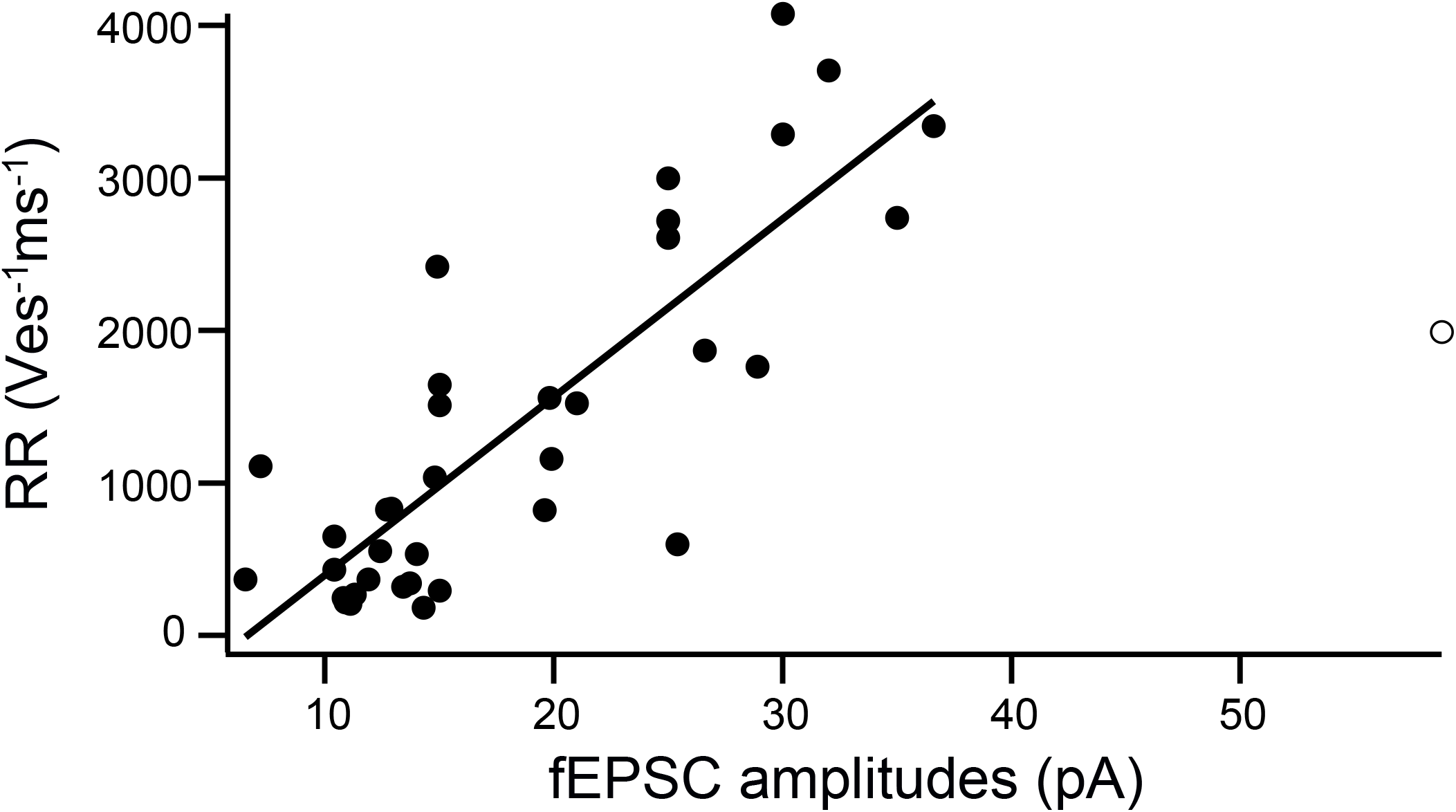
Correlation between peak RR and fEPSC amplitudes. Peak release rates (RR) were plotted against the corresponding fEPSC amplitudes. The regression line indicates a linear correlation (P_r_ = 0.83).

**Supplementary Fig. 5.**
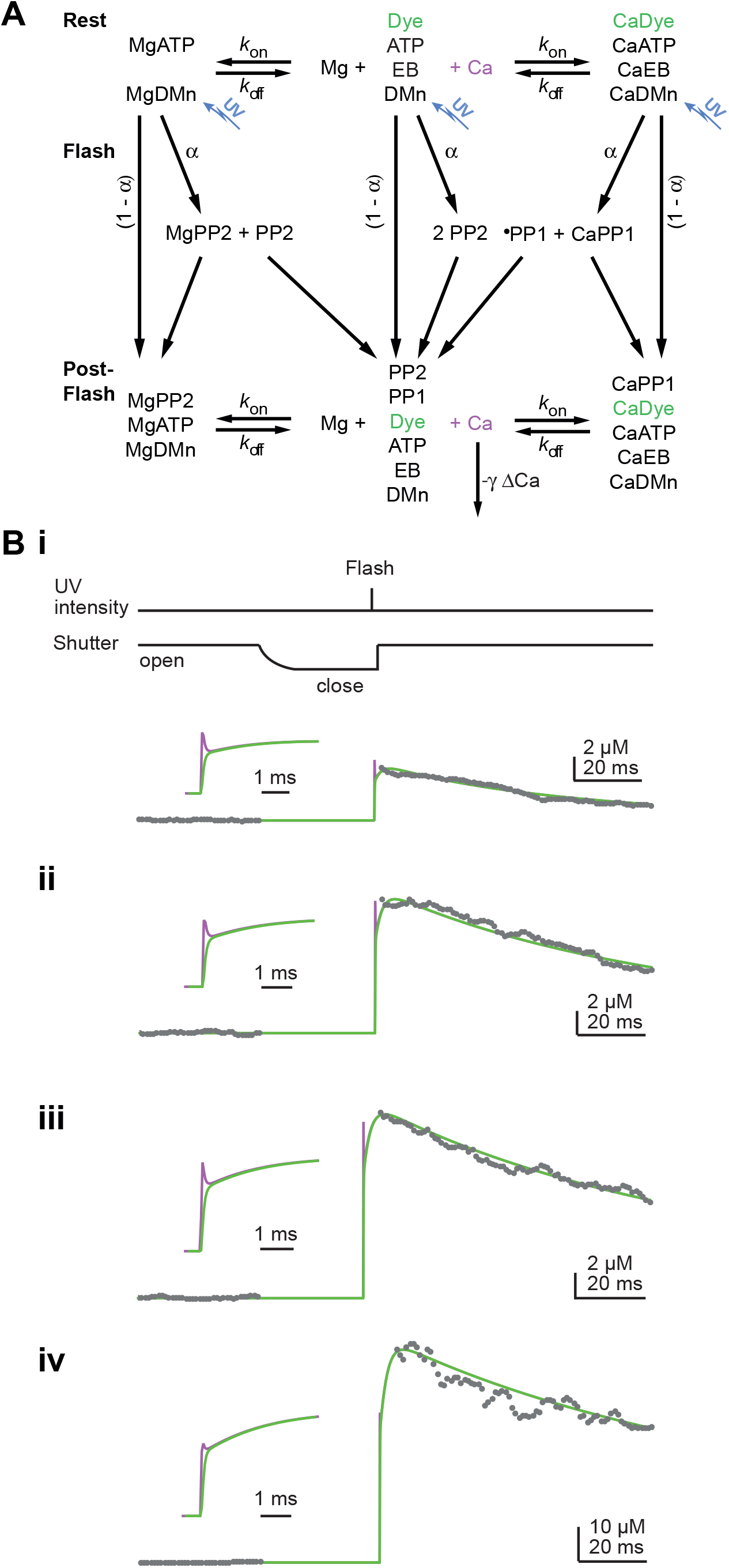
Model of flash-evoked Ca^2+^ elevations. **(A)** Scheme of the chemical reactions that were implemented in the model of flash-induced Ca^2+^ uncaging. The model covered Ca^2+^ (Ca) and Mg^2+^ (Mg)-binding to the indicator dye (OGB5N), to DM-nitrophen (DMn) and to buffers (ATP and an endogenous buffer (EB)). The forward (*k*_on_) and backward (*k*_off_) rate constants differ between chemical species. Upon simulated UV flash photolysis, a fraction (α) of metal bound and free DMn made a transition to different photoproducts (PP1 and PP2) (*25, 31*). The model parameters are given in **Supplementary Table 1**. (**Bi**) Drawings of the laser intensity profile and the shutter action. The shutter velocity was slow during closing but rapid during opening. (**Bii-iv**) Measured elevations in [Ca^2+^]_i_ (grey dots) induced by photolysis from CaDMn with different UV laser intensities and/or durations. Superimposed on the data are Δ[Ca^2+^]_i_ (purple line) and Δ[Ca^2+^]_i_ as reported by OGB5N (green line) simulated with the model in (A) by increasing the parameter α from (ii) to (iv). After ∼0.5 ms the dye reliably reflects the time course of [Ca^2+^]_i_. Note that under our recording conditions a predicted initial spike-like overshoot in Δ[Ca^2+^]_i_ is very brief and not prominent.

**Supplementary Fig. 6.**
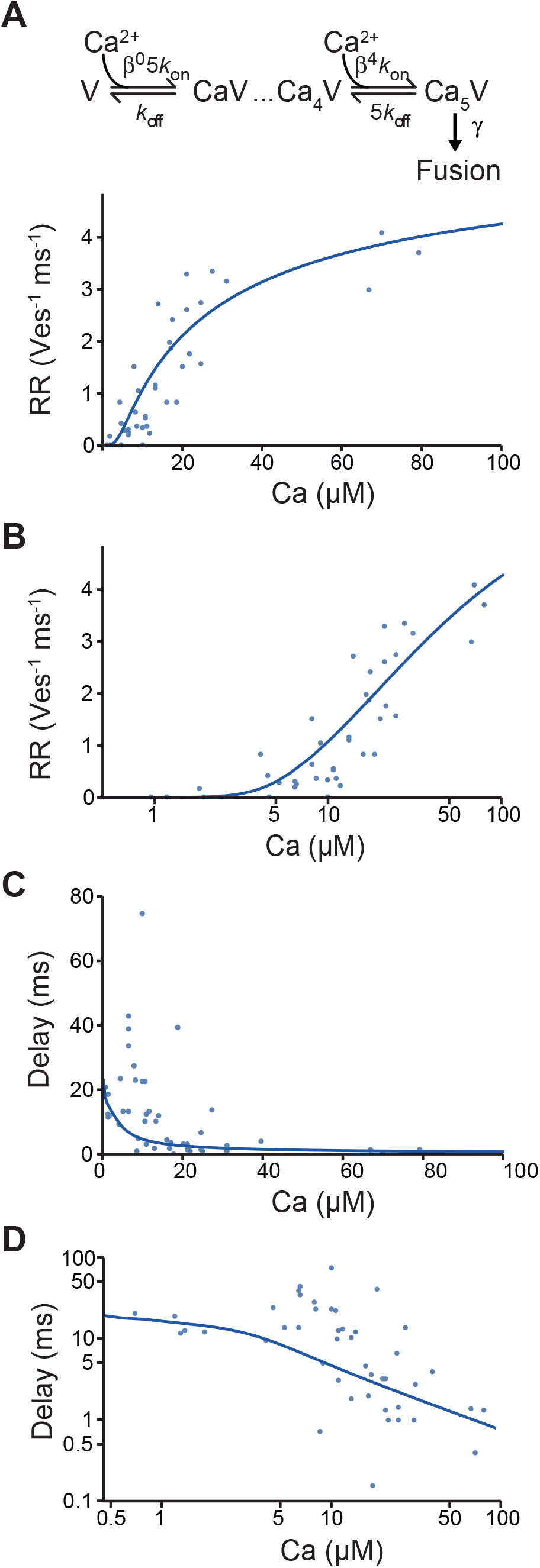
Model of the Ca^2+^-dependency of Syt1-triggered release. **(A)** *Top*: Reaction scheme of the five-site model used to simulate the data of flash-evoked release. Δ realizes positive cooperativity in the forward binding reactions. *Bottom*: Data from **Fig. 5** (circles) that were fitted by the model (curve). **(B)** As in (A, *bottom*) but with a logarithmic x-axis. **(C)** As in (A, *bottom*) but for the synaptic delays. **(D)** As in (C) but on a double-logarithmic scale.

**Supplementary Fig. 7.**
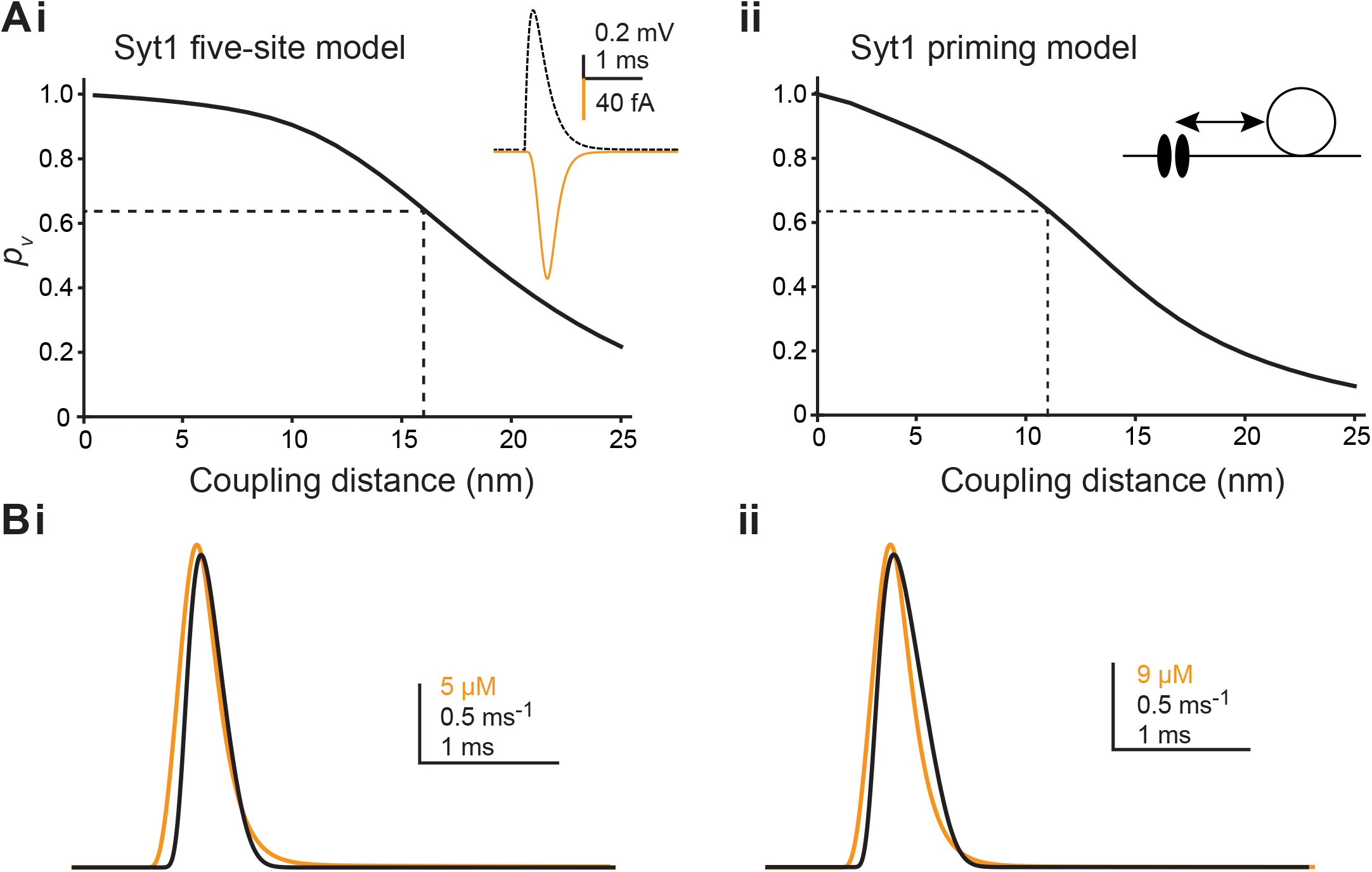
Estimating coupling distance and AP-mediated Ca^2+^ at Syt1. The coupling distance between a single Ca_v_2.1 channel and the release sensor was estimated based on AP-mediated Ca^2+^ signals measured in boutons of mature L5PNs, the measured vesicular release probability (*p*_v_ = 0.63), and spatially resolved simulations as described previously (*11*), except that the sensor model was replaced by the newly established ‘Syt1 models’. (**Ai**) Simulated *p*_v_ at increasing coupling distance using the Syt1 five-site model (**Supplementary Fig. 6**). The dashed line indicates the coupling distance at which the experimentally determined *p*_v_ of 0.63 was predicted. *Inset*: AP waveform (black) and Ca^2+^ influx through a single Ca_v_2.1 channel (orange). (**Aii**) As in (i) but for the Syt1 priming model (**Fig. 5**). (**Bi**,**ii**) [Ca^2+^]_i_ transients (orange) and time courses of the release rates (black) at the coupling distances determined in (A) with the five-site model (i) or the priming model (ii).

**Supplementary Fig. 8.**
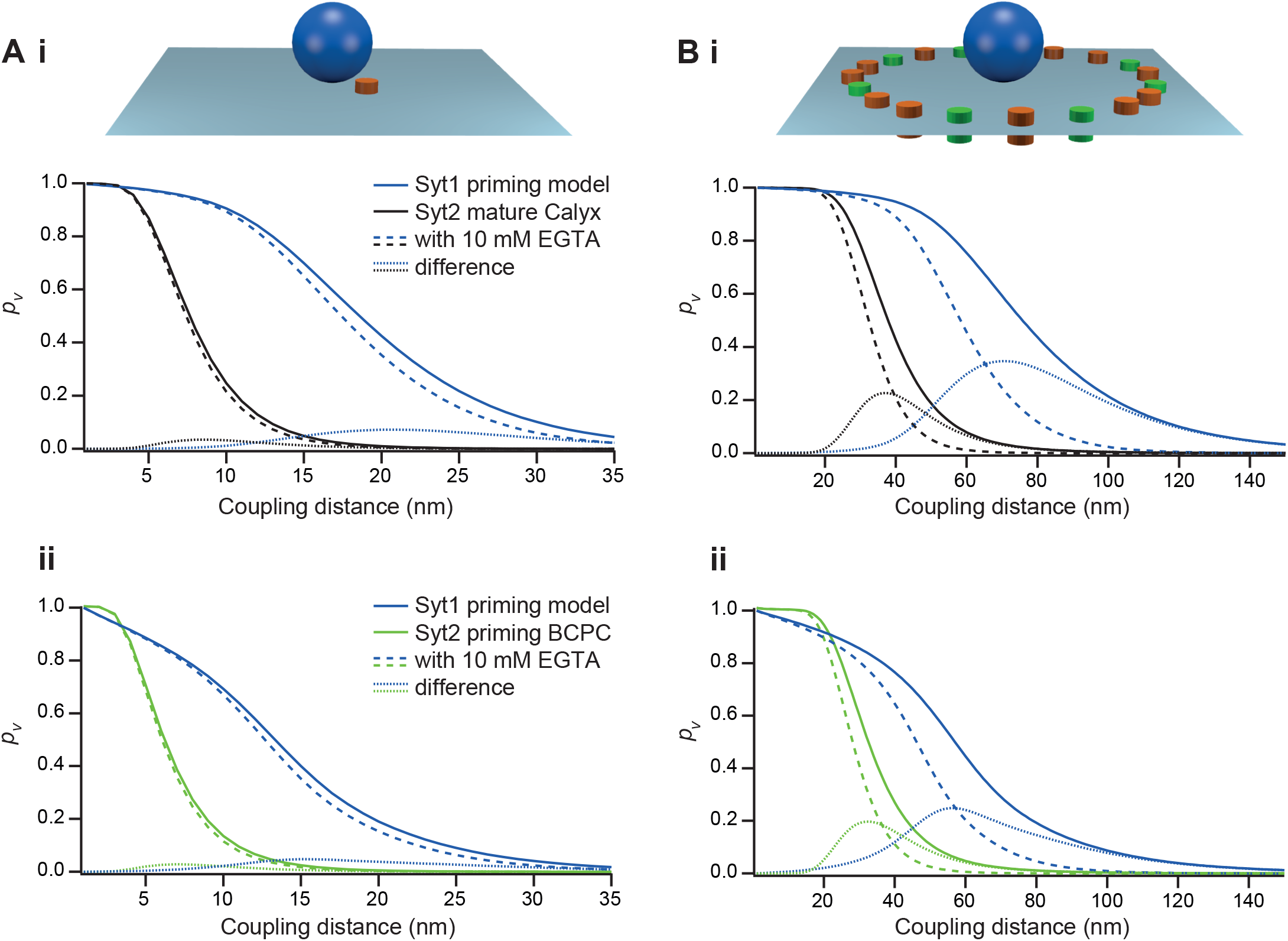
Comparison of Syt1- and Syt2-triggered release with different presynaptic nanotopographies. (**Ai**) Simulated *p*_v_ at increasing coupling distance in a nanodomain coupling regime between a single Ca_v_2.1 channel and the release sensor (inset). An AP-mediated presynaptic [Ca^2+^]_i_ signal was used as described in ref.14 (see **Supplementary Fig. 7**). The release sensor was simulated either by the Syt1 five-site model (blue) or a Syt2 five site model of the mature calyx of Held (black) (*22*). The dashed lines are simulations in the presence of the slow Ca^2+^ buffer EGTA (10 mM). The doted lines represent the difference between control simulations and simulations in the presence of EGTA. (**Aii**) Same as in (i) but for the Syt1 priming model (blue) in comparison to a Syt2 priming model from an inhibitory synapse (*6*). (**Bi**) As in (Ai) but in a microdomain coupling regime with 7 Ca_v_2.1 (brown) and 5 Ca_v_2.2 (green) channels (*11*) (inset). (**Bii**) As in (Aii) but for the microdomain as in (Bi).

## Notes

### Competing Interest Statement

The authors have declared no competing interest.

### Summary of Updates

Abstract, Introduction and Discussion were shortened

